# *Staphylococcus aureus* staphyloxanthin expression is not controlled by Hfq

**DOI:** 10.1101/2019.12.17.876839

**Authors:** Wenfeng Liu, Pierre Boudry, Chantal Bohn, Philippe Bouloc

## Abstract

**Objective:** The golden color of *Staphylococcus aureus* is due to the synthesis of carotenoid pigments. In Gram-negative bacteria, Hfq is a global posttranscriptional regulator, but its function in *S. aureus* remains obscure. The absence of Hfq in *S. aureus* was reported to correlate with production of carotenoid pigment leading to the conclusion that Hfq was a negative regulator of the yellow color. However, we reported the construction of *hfq* mutants in several *S. aureus* strains and never noticed any color change; we therefore revisited the question of Hfq implication in *S. aureus* pigmentation.

**Results:** The absence or accumulation of Hfq does not affect *S. aureus* pigmentation.

## INTRODUCTION

*Staphylococcus aureus* is a major pathogen responsible for numerous diseases from minor skin infection to septicemia, affecting humans and other animals. Its name *“aureus”* comes from the golden color of strains that express carotenoid pigments [1]. These pigments contribute to oxidative stress and neutrophil resistance, and virulence [2]. The carotenoid biosynthetic operon (*crtMNOPQ*) leading to the synthesis of staphyloxanthin regulated by σ^B^ [3, 4], an alternative σ factor that also controls a large number of general stress genes. σ^B^ activity depends on RsbU, its positive regulator [5, 6]. Numerous strains, including the *S. aureus* model NCTC8325, have *rsbU* mutations that prevent σ^B^ activity and *crt* operon expression, such that colonies are white. In addition, mutations in 37 genes were show to result in the loss of a yellow pigmentation [5, 7].

Hfq is an RNA chaperone needed for activity of numerous regulatory RNAs in Gram-negative bacteria [8]. However, its role in Gram-positive bacteria, with the exception of *Clostridium difficile* [9], remains enigmatic [10]. Hfq functionality from different species is often tested by interspecies complementation tests. However, expression of *hfq* genes from Gram-positive bacteria *S. aureus* and *Bacillus subtilis* in *Salmonella* could not compensate the absence of endogenous *hfq*, indicating a functional difference between Gram positive and negative Hfq [11, 12].

We previously compared phenotypes of *S. aureus hfq* mutants with their isogenic parental strains and observed no detectable difference associated with the absence of Hfq in the tested conditions [13]. However, our results were partly challenged by a publication reporting that carotenoid pigment production was increased in *hfq*-negative strains [14]. Here we use nine different *S. aureus* strains to show that Hfq absence or overexpression has no effect on pigment expression.

## MAIN TEXT

### Methods

#### Bacterial strains, plasmids and growth conditions

Bacterial strains, plasmids and primers used in this study are listed in Table 1. Allelic replacements of *hfq*^+^ by Δ*hfq::cat* were either performed by □11-phage mediated transduction using RN4220 *hfq*::cat as a donor strain or by homologous recombination using pMADΔhfq::cat [13, 15]. The Δ*hfq::cat* deletion in SAPHB5 was verified by Southern blot and subsequent Δ*hfq::cat* transductants were verified by PCR as described [13].

**Table 1:**
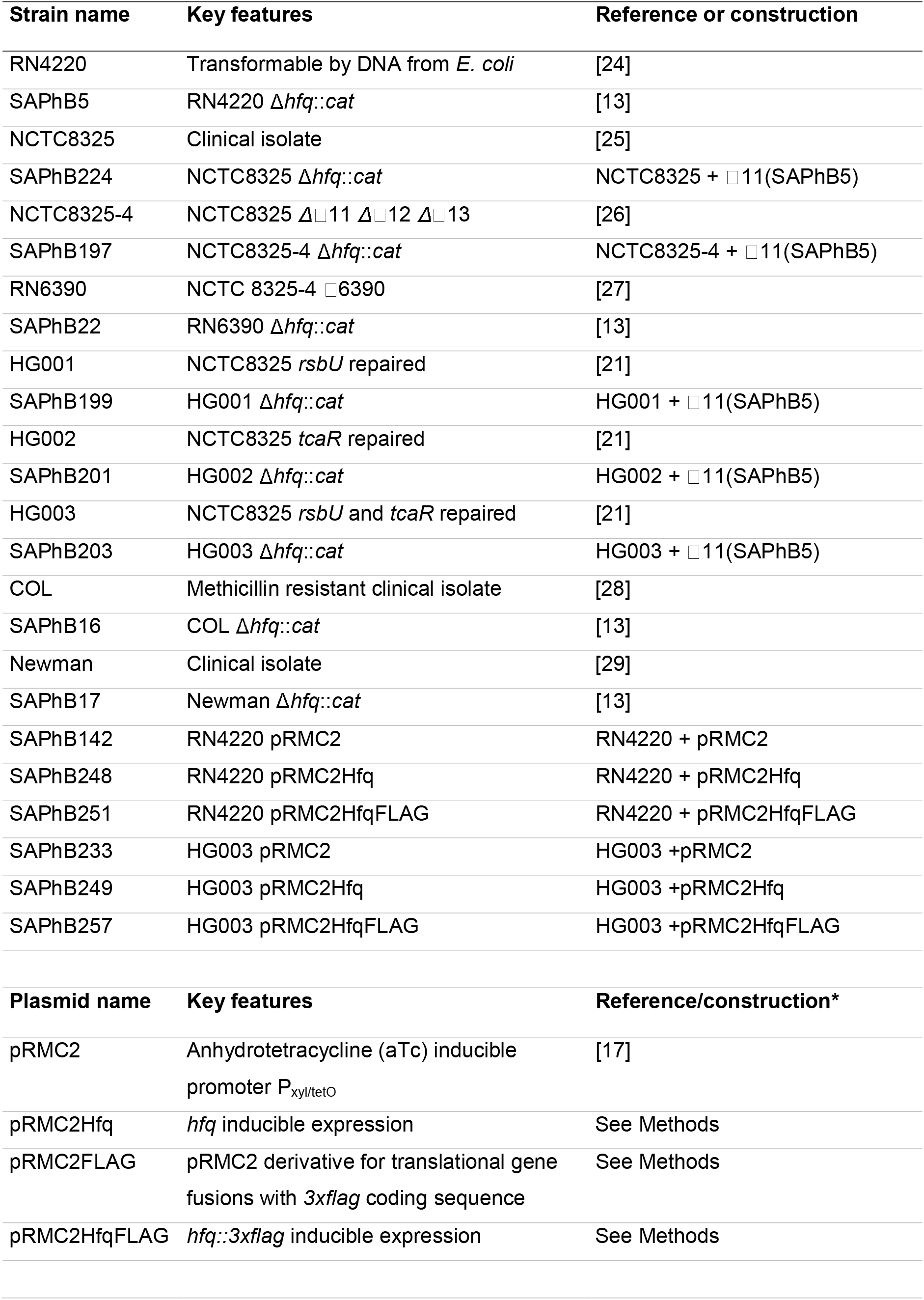
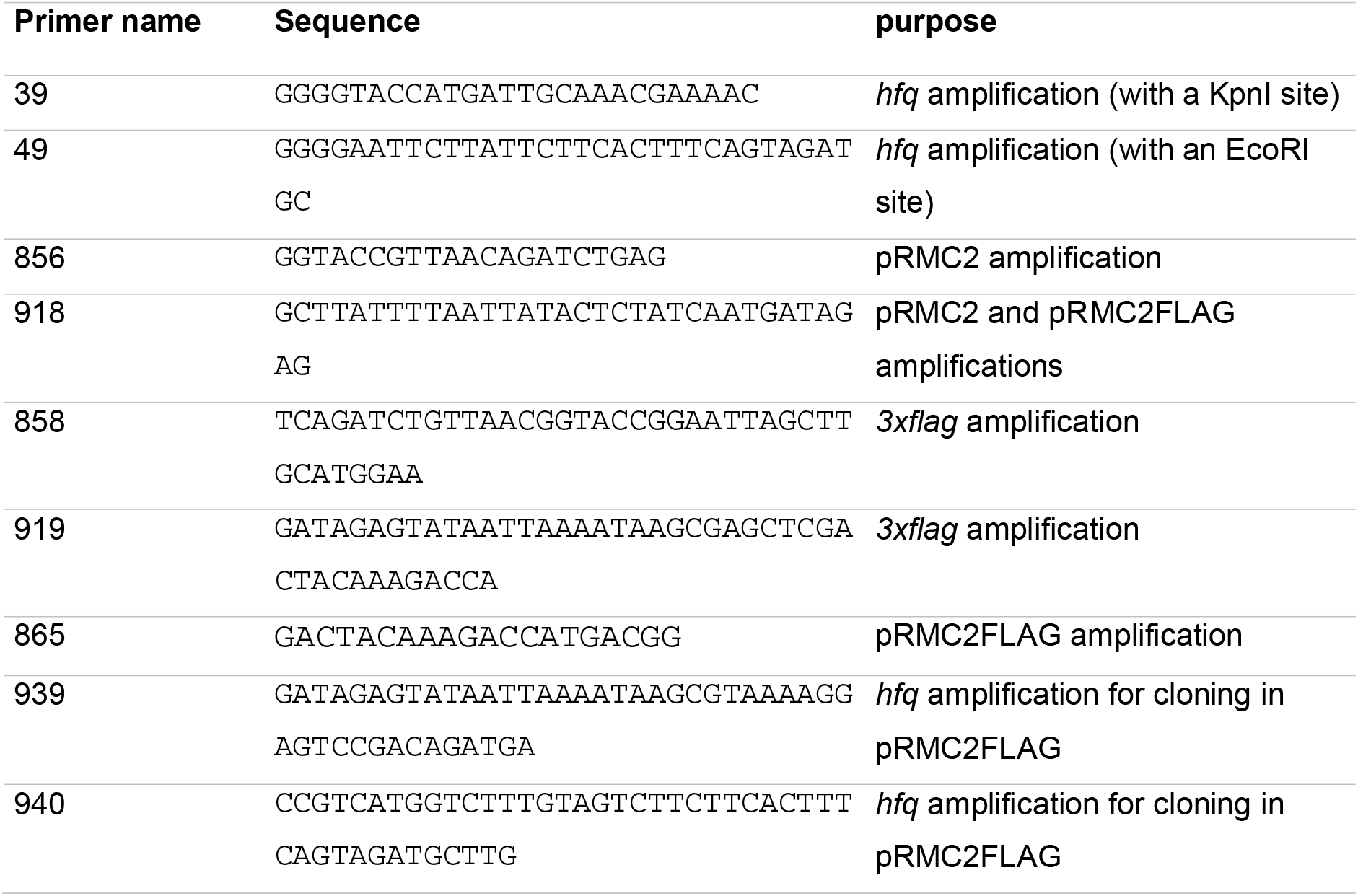
*Staphylococcus aureus* strains, plasmids and primer used for this study.

Engineered plasmids were constructed as described [16]. Conditional *hfq* expression was obtained by cloning *hfq* under the xyl/tetO promoter in pRMC2 [17] and pRMC2FLAG (Table 1). pRMC2Hfq allowing *hfq* conditional expression was obtained as follows: pRMC2 and PCR-amplified *hfq* (using primers 39/49 on HG003 DNA) were KpnI-EcoRI digested and ligated together. pRMC2FLAG was engineered for conditional expression of 3xFLAG-tagged proteins as followed: pRMC2 and pSUB11 [18] were PCR-amplified using primers 856/918 and 858/919, respectively. The two resulting products, *i.e.* pRMC2 and *3xflag* coding sequence, were assembled using the Gibson method [19]. pRMC2HfqFLAG, allowing conditional expression of Hfq::3xFLAG, was obtained as follows: pRMC2FLAG and *hfq* HG003 were PCR-amplified using primers 918/865 and 939/940, respectively. The two resulting products were assembled using the Gibson method.

Bacteria were grown in BHI medium at 37°C. For strains containing pRMC2 and derivatives, chloramphenicol 5 μg.ml^-1^ was added to media. Expression from pRMC2 and derivatives was achieved by anhydrotetracycline (aTc) 250 ng.ml^-1^ addition to growth media.

#### Protein extraction and Western blotting

Overnight cultures were diluted 1000 times in fresh medium. After 3h, aTc was added. 10 min and 30min later, cells were harvested by centrifugation (16.000 g for 2 min), resuspended in 400μl Tris HCl buffer (50 mM, pH 6.8) and lysed using a FastPrep (3 cycles of 45 sec at 6.5 m.s^-1^). Cell debris was removed by centrifugation (16.000 g for 10 min). Protein concentration was determined by Bradford assays [20]. For each sample, 3 μg of protein extract was separated on a polyacrylamide gel (Blot™ 4-12% Bis-Tris Plus, Invitrogen). After electrophoresis, proteins were transferred to a polyvinylidene fluoride membrane (iBlot 2 PVDF Mini Stacks, Invitrogen). For blotting and washing, an iBind™ Flex Western System was used according to supplier’s instructions. Membranes were probed with the primary polyclonal ANTI-FLAG antibody produced in rabbit (Sigma) at a 1/15,000 dilution. A rabbit secondary antibody conjugated to horseradish peroxidase (Advansta) was used at a 1/25,000 dilution. Bioluminescent signal was detected with the WesternBright™ ECL-spray (Advansta) using a digital camera (ImageQuant™ 350, GE Healthcare).

### Results

#### The absence of Hfq does not alter *S. aureus* pigmentation

In 2010, Liu *et al.* reported that “deletion of *hfq* gene in *S. aureus* 8325-4 can increase the surface carotenoid pigments” [14]. Their work was performed using an allele called Δ*hfq-8325* in which the *hfq* coding sequence was replaced by a kanamycin cassette. The *hfq* chromosomal deletion was constructed in strain RN4220 and then transduced into NCTC8325-4, RN6390, COL and ATCC25923 by phage □11. We constructed a similar *hfq* deletion in RN4220, except that the *hfq* coding sequence was replaced by a chloramphenicol resistant gene (Δ*hfq*::cat); this allele was transduced into RN6390, COL and Newman by □ 11-phage mediated transduction [13]. Note that RN4220, RN6390 and COL strains were used for both studies. As we did not notice a change of color when the Δ*hfq*::cat allele was introduced into these strains, this information was not reported [13]. In view of the previous report, we focused this work on the possibility that Hfq could affect *S. aureus* pigment expression.

NCTC8325 isolated in 1960 from a sepsis patient is the progenitor of numerous strains including NCTC8325-4 (cured of three prophages) which itself gave RN6390 and RN4220 [21]. As these descendants were mutagenized, they carry several mutations that may affect their phenotypes. NCTC8325 has a deletion of 11 bp in *rsbU* and a point mutation in *tcaR.* The derivatives HG001 *(rsbU* restored), HG002 *(tcaR* restored), HG003 *(rsbU* and *tcaR* restored) were constructed to perform physiological studies in a non-mutagenized background [21]. All these strains derived from NCTC8325, except HG001 and HG003 (which have a functional σ^B^ factor), give rise to white colonies (Figure 1). In addition to those reported [13], we constructed Δ*hfq*::cat derivatives in NCTC8325, NCTC8325-4, HG001, HG002 and HG003 (Table 1). In contrast to results reported in Liu *et al.*, deletion of the *hfq* gene in all tested strain backgrounds had no effect on pigmentation (Figure 1A). Note that COL, Newman are not NCTC8325 derivatives.

**Fig 1:**
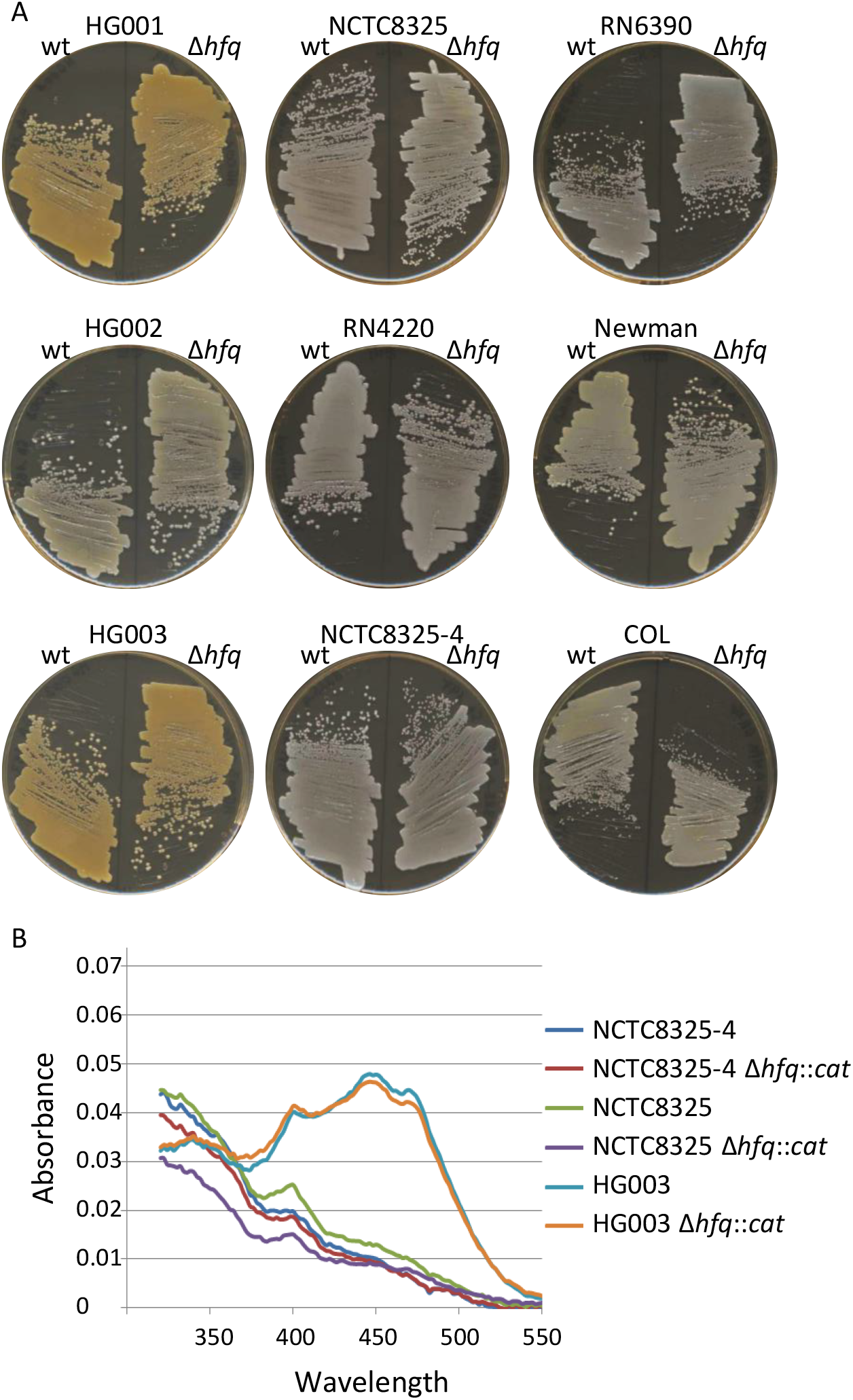
Absence of Hfq does not affect *S. aureus* pigmentation. The indicated strains were grown overnight in BHI and then A) streaked on BHI agar or B) assayed for spectral profiles as described [1].

Spectral profiles highlighting *S. aureus* carotenoid production were determined as described [1] for three strains and their *hfq* derivatives after growth for 24h in BHI. HG003 and HG003 Δ*hfq*::cat gave equivalent profiles with three pics characteristic of carotenoid production. In contrast, NCTC8325-4 and RN1 had spectra characteristic of no or very little carotenoid production. As expected from our visual observation (Figure 1A), the spectra of Δ*hfq* derivatives did not differ from those of their respective parental strains (Figure 1B).

#### Hfq overexpression does not alter *S. aureus* pigmentation

In the above-described strains, *hfq* is possibly poorly expressed, in which case *hfq* deletions would not lead to detectable phenotypes. We therefore tested the effects of an inducible Hfq expression system on pigment production. If the absence of Hfq leads to yellow colonies as proposed [14], the presence of Hfq could lower pigment production and lead to white colonies. To address this point, *hfq* was cloned under the control of the P_xyl/tetO_ promoter in multi-copy plasmid pRMC2 [17] leading to pRMC2Hfq. *hfq* expression in strains harboring pRMC2Hfq was induced upon aTc addition to media. To confirm that P_xyl/tetO_ was effectively driving *hfq* expression, a pRMC2Hfq derivative was engineered harboring a *3xflag* sequence inserted in frame at the end of the *hfq* open reading frame. The resulting plasmid, pRMC2HfqFLAG is a proxy for expression from pRMC2Hfq. HG003 was transformed with pRMC2, pRMC2Hfq and pRMC2HfqFLAG. The protein Hfq::3xFLAG was detected upon aTc induction by western blotting using FLAG antibodies (Figure 2A). We inferred from this result that addition of aTc to strains harboring pRMC2Hfq lead to Hfq synthesis. The RN4220 white and HG003 yellow colors were not affected by the presence of either pRMC2, pRMC2Hfq or pRMC2HfqFLAG and remained identical upon aTc addition to growth medium (Figure 2B).

**Fig 2:**
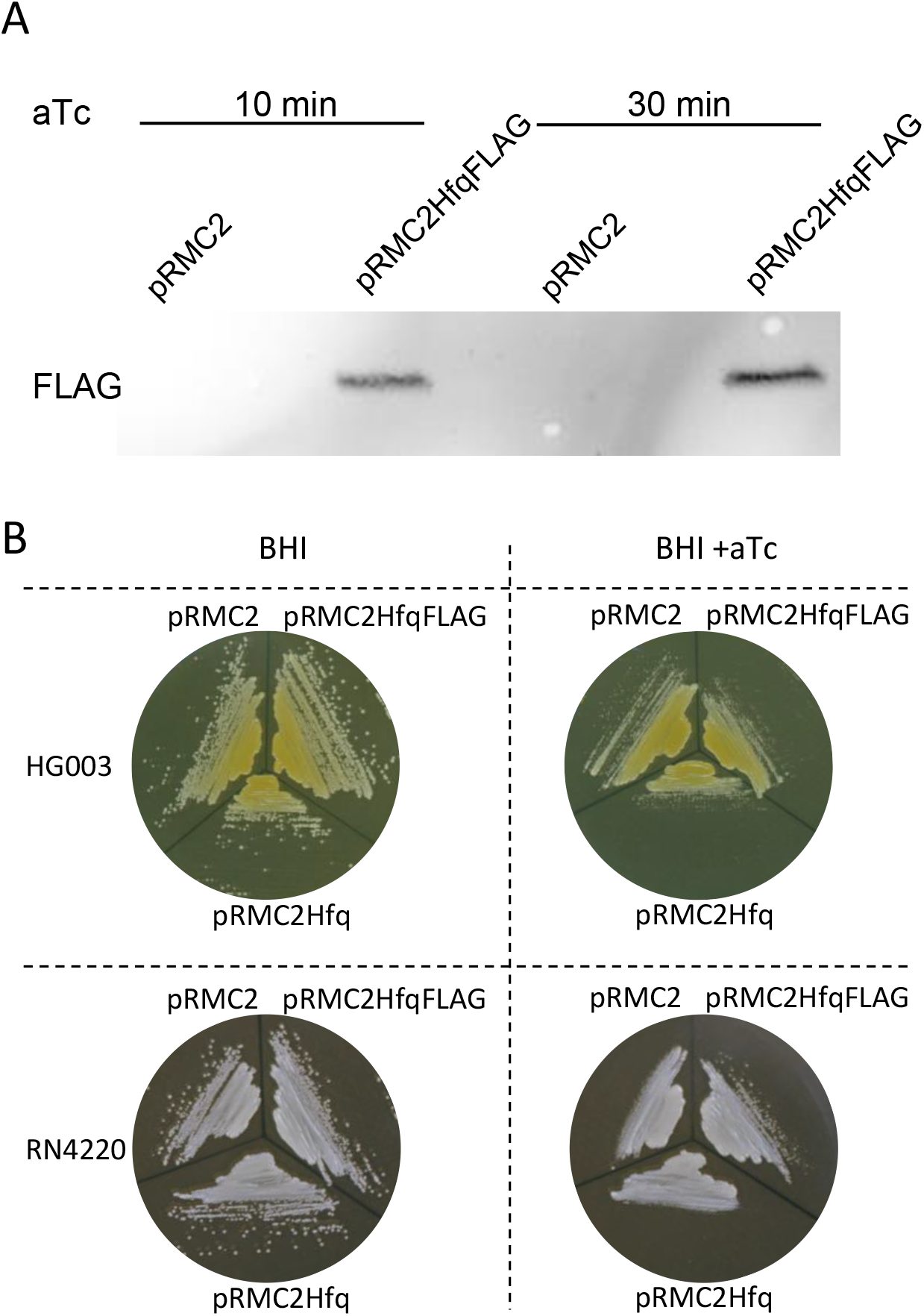
Accumulation of Hfq does not affect *S. aureus* pigmentation. The indicated strains were grown in BHI supplemented with chloramphenicol. They were then A) assayed by Western blotting for *hfq::3xflag* expression (see Methods) and B) streaked on BHI agar supplemented with aTc when indicated.

### Conclusion

Our results show that neither the absence, nor the accumulation of Hfq affects pigmentation of *S. aureus:* Hfq does not appear to regulate staphyloxanthin synthesis. Our conclusions are supported by E.J. Tarrant PhD dissertation showing an NCTC8325 *hfq* mutant that remained unpigmented [22]. Of note, *Pseudomonas aeruginosa* reportedly induces pigment production of *S. aureus*, however, this effect was independent of *hfq* transcription [23]. In addition, color variation in USA300 strain was screened in a genome-wide transposon mutant library, and the *hfq* inactivation was not reported to affect *S. aureus* pigmentation [7].

While the *hfq* gene is absent in some Firmicutes *(e.g.* Lactobacillales), it is conserved in all *S. aureus*, suggesting that it plays a crucial function, however not related to pigment expression. The quest to find the Staphylococcal Hfq function remains open.

## LIMITATIONS

Our conclusion is in contradiction with Liu *et al.* results concerning the effect of Hfq on *S. aureus* pigmentation [14]. We cannot rule out that our observation is limited to specific *S. aureus* strains. However, we used an NCTC8325-4 *hfq* derivative similar the one used in the previous study. Furthermore, the present results are strengthened by the construction of *hfq* mutants in numerous *S. aureus* backgrounds. The discrepancy between our and Liu *et al.* 2010 [14] results, is a possible inadvertent selection of mutants with altered color patterns (as shown in [22]) in the former study.

## List of abbreviations

PCR: polymerase chain reaction
BHI: brain heart infusion
aTc: anhydrotetracycline

## DECLARATIONS

### Ethics approval and consent to participate

Not applicable

### Consent for publication

Not applicable

### Availability of data and materials

All data generated or analyzed during this study are included in this published article. Strains and plasmids are available from the corresponding author on reasonable request.

### Authors’ contributions

PhB designed the experiments and wrote the manuscript. WL, PiB and CB performed the experiments, analyzed data and revised the manuscript. All authors read and approved the final manuscript.

## Acknowledgements

We thank Sandy Gruss, Annick Jacq and Yan Jing for critical reading of the manuscript, helpful discussions and warm support.

